# Atheroma plaque microenvironment stimulates kynurenine production by macrophages to induce endothelial adhesion molecules in the context of atherogenesis

**DOI:** 10.1101/2023.07.19.549799

**Authors:** Kevin Van Dongen, Damien Leleu, Thomas Pilot, Antoine Jalil, Léa Mangin, Louise Ménégaut, Audrey Geissler, Stoyan Ivanov, Aline Laubriet, Valentin Crespy, Maxime Nguyen, Eric Steinmetz, David Masson, Charles Thomas, Thomas Gautier

## Abstract

Cardiovascular diseases, including atherosclerosis, are major causes of morbidity and mortality worldwide. Here, we investigate the role of the kynurenine pathway (KP) in macrophages in the context of atheroma plaque microenvironment and its impact on atherogenesis. Using an in vitro model of primary human macrophages, we observed that exposure to plaque homogenates induces a marked increase in the early steps of the KP which impacts on kynurenine production. This was confirmed by immunostaining on human plaque of carotid arteries. Further investigation into the underlying molecular mechanisms revealed that LXR signaling contributes to this plaque microenvironment-induced KP activation. We showed that kynurenine released from macrophages affected endothelial cells, leading to increased expression of ICAM-1 and VCAM-1 in an AhR-dependent manner. Consistently with the proatherogenic effects, in a cohort of atherosclerotic patients, we observed higher levels of plasma kynurenine in patients with lower extremity arterial disease. In line with the results of in vitro investigations, the plasma kynurenine levels were associated plaque oxysterol content. Using a multiple logistic regression model, we showed that plasma kynurenine was independently associated with lower extremity arterial disease in atherosclerotic patients. Altogether, our data indicate that the activation of KP in macrophages in the context of atheroma plaque is partially mediated by LXR axis and leads to the release of kynurenine. This, in turn, contributes to the exacerbation of both local and peripheral atherosclerosis particularly through the activation of endothelial cells.

## Introduction

Atherosclerosis represents a persistent inflammatory disorder characterized by the gradual buildup of lipids and immune cells, particularly macrophages, within the arterial intima. This pathological process ultimately leads to the formation of atherosclerotic plaques, which can precipitate abrupt cardiovascular events such as myocardial infarction or stroke through the rupture of vulnerable plaques. Consequently, there arises a pressing need for novel therapeutic strategies aimed at treating atherosclerosis or impeding its progression.

Monocytes and macrophages assume pivotal roles throughout all stages of atheroma plaque formation, encompassing its initiation, progression, and rupture. Of particular significance is their involvement in instigating and perpetuating a chronic inflammatory response, which constitutes a fundamental mechanism driving the pathophysiology of atherosclerosis and the occurrence of acute cardiovascular events ^1^.

The efficacy of interventions designed to selectively target the inflammatory response, thus effectively mitigating the occurrence of cardiovascular events associated with atherosclerosis has been recently demonstrated ^2^. However, our understanding of the precise mechanism underlying the recruitment and subsequent activation of monocytes/macrophages within the atheroma plaque microenvironment remains incomplete. This complexity arises in part due to the utilization of animal models, which only partially recapitulate the pathophysiology of atherosclerosis and have limited capacity to investigate cellular-level mechanisms.

Our team recently developed an in vitro model of primary human macrophages exposed to atherosclerotic plaque from patients who underwent endarterectomy ^3, 4^. We have been able to demonstrate that this model reproduces the profile of inflammatory macrophages found within the human atherosclerotic plaque, including a significant reprogramming of the expression of genes encoding for inflammatory cytokines, metalloproteinases or lipid metabolism ^3, 4^. Using this model, we highlighted the ability of atheroma plaque homogenates to activate IL-1β in a Liver-X-receptors (LXR)-dependent manner as well as the co-localization of LXR with HIF-1α and IL-1 β in human macrophage-rich regions of atheroma plaques. By contrast, as previously reported^5^, LXR activation was associated to a reduction of IL-1β expression in murine macrophages^4^. LXRs represent an essential link between cholesterol accumulation and inflammatory response in macrophages^6^. These nuclear receptors are activated by cholesterol derivatives such as oxysterols and desmosterol, compounds that accumulate in cells after cholesterol loading ^7, 8^. In turn, LXRs activate genes that limit cholesterol accumulation by stimulating cholesterol efflux and inhibiting cholesterol uptake. As we, and others, showed, LXRs also modulate the inflammatory response of macrophages by several distinct mechanisms^5, 9, 10^, namely through LXR/HIF-1α signaling^4^. Importantly, our results demonstrate that the LXR/HIF-1α axis is not restricted to IL-1β. Indeed, other HIF-1α-dependent metabolic pathways are significantly activated by LXR and display similar human selectivity. Genes involved in glycolysis such as GLUT1 and HK2 ^4^ strongly induced by LXR in a HIF-1α-dependent manner in human macrophages, whereas LXR activation has almost no impact in murine macrophages. More recently, we further demonstrated that conditioning of human macrophages by plaque homogenates induces expression of several genes involved in glucose uptake and glycolysis including glucose transporter 1 (SLC2A1) and hexokinases 2 and 3 (HK2 and HK3). We showed that this activation was significantly correlated to the oxysterol content of the plaque samples and was associated with a significant increase in the glycolytic activity of the cells. Moreover, we highlighted that pharmacological inverse agonist of the oxysterol receptor liver X receptor (LXR) partially reverses the induction of glycolysis genes without affecting macrophage glycolytic activity.

It is of utmost significance to highlight a fundamental distinction between human and murine macrophages with regards to their metabolic response upon activation of Liver X Receptors (LXRs). Our research demonstrates that LXR agonists elicit divergent metabolic shifts in human and murine macrophages, wherein human macrophages experience a prominent induction of glycolysis following LXR activation. It has been established that certain macrophage subpopulations within atheroma plaques exhibit heightened glucose uptake activity. Nonetheless, the precise molecular mechanisms and pathophysiological implications of this heightened glucose influx remain elusive. Our findings provide novel insights into these aspects and propose a potential involvement of LXRs and their activators, particularly cholesterol/oxysterols accumulation, in regulating the metabolic activity of macrophages within atheroma plaques. Furthermore, it is highly probable that the metabolic reprogramming of human plaque macrophages extends beyond glucose metabolism. Consequently, in our present study, we sought to explore these aspects further by employing a similar approach utilizing human primary macrophages exposed to homogenates derived from patient plaques. Our investigations revealed a substantial alteration in the kynurenine pathway due to the plaque microenvironment, and we identified several crucial genes associated with this pathway, such as kynurenine monooxygenase (KMO), which were regulated by the LXR/HIF-1α axis. Additionally, we demonstrated that the subsequent production of kynurenine led to increased expression of adhesion molecules on human endothelial cells. Interestingly, in a cohort of patients who underwent endarterectomy, we observed a correlation between elevated plasma kynurenine levels and the presence of lower limb atherosclerosis.

## Results

### LXR signaling contributes to plaque microenvironnement-induced kynurenine pathway

Our team recently developed an *in vitro* model of primary human macrophages exposed to atherosclerotic plaque homogenates from patients who underwent endarterectomy^3, 4^. Using a similar strategy combined to RNA sequencing, we observed an upregulation of the kynurenine pathway (KP) in primary human macrophages exposed to plaque microenvironment (Fig. 1A-B). The KP plays a central metabolic role in the cell as it constitutes one of the pathways for NAD^+^ regeneration, along with the Preiss-Handler pathway and the salvage pathway^11^. This pathway has long been considered a minor contributor to intracellular NAD^+^ regeneration, except in the liver. However, recent studies have shown that the kynurenine pathway significantly contributes to NAD^+^ production in macrophages^12^. Interestingly, we showed here that exposure to plaques homogenates induces a marked increase in the early steps of KP. By contrast, gene expression of *quinolinate phosphoribosyltransferase* (QPRT) and of genes of Preiss-Handler pathway and the salvage pathway were downregulated or unchanged (Fig 1B-C, Fig. S1). Notably, a concomitant decrease in intracellular levels of NAD^+^ was observed in macrophages exposed to plaque homogenates (data not shown). Consequently, it is reasonable to propose that the effects resulting from the upregulation of the initial steps of the KP predominantly impact the production and release of KP metabolites rather than exerting a significant influence on the bioenergetic processes of macrophages. Prior to delving into a comprehensive investigation of these aspects, we explored of the underlying molecular mechanisms involved.

**Figure 1:**
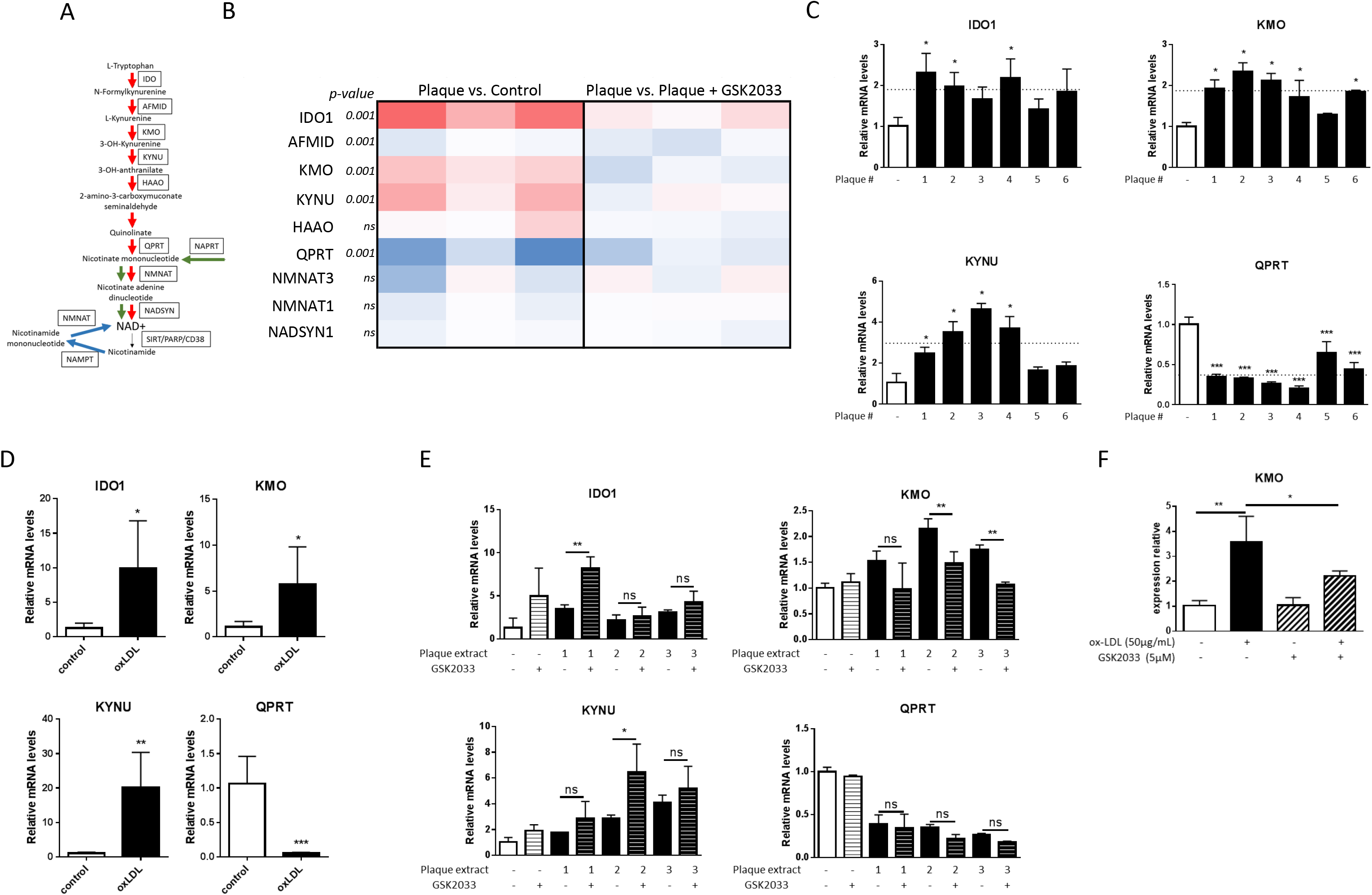
LXR signaling contributes to atheroma plaque microenvironnemen-dependent regulation of kynurenine pathway in human macrophages. A. Schematic view of NAD^+^ synthesis pathway. In red, de novo pathway (kynurenine pathway). In green, Preiss-Handler pathway, In blue, salvage pathway. B. Gene expression heatmap of key enzymes of the kynurenine pathway. Differential gene expression determined by RNA sequencing in primary human macrophages exposed for 48 h with patient atheroma plaque homogenates (left, fold change compared to vehicle (DMSO)) or patient atheroma plaque homogenates plus the LXR antagonist GSK2033 (1μM) (right, fold change, atheroma plaque versus atheroma plaque plaque plus GSK2033). n=3. P value determined by Student’s t-test. C: Relative mRNA levels of IDO1, KMO, KYNU and QPRT in primary human macrophages exposed for 48h to patient atheroma plaque homogenates. The dotted line indicates average change in gene expression obtained with the different atheroma plaques homogenates. Data are representative of 4 independent experiments with different healthy donors. Bar graphs represent mean ± s.d * p<0.05; ** p<0.01; *** p<0.001 determined by Student’s *t*-test. D. Relative mRNA levels of IDO1, KMO, KYNU and QPRT in primary human macrophages (n=5 healthy donors) treated for 24h with oxLDL (50µg/mL). Data are representative of 3 independent experiments with different healthy donors. Bar graphs represent mean ± s.d * p<0.05; ** p<0.01; *** p<0.001 determined by Student’s *t*-test. E. Relative mRNA levels of IDO1, KMO, KYNU and QPRT in primary human macrophages exposed for 48h with patient atheroma plaque homogenates alone or in combination with the LXR antagonist GSK2033 (1µM). Data are representative of 4 independent experiments with different human healthy donors. Bar graphs represent mean ± s.d. * p<0.05; ** p<0.01; *** p<0.001 determined by Student’s *t*-test. F. Relative mRNA levels of KMO in primary human macrophages exposed for 24h to oxLDL alone or in combination with the LXR antagonist GSK2033 (5µM). Bar graphs represent mean ± s.d. * p<0.05; ** p<0.01; *** p<0.001 determined by one way ANOVA corrected for multiple comparison.

First, we showed that exposure of primary human macrophages to oxidized-LDLs (oxLDLs) leads to the same gene expression profile with regards of *IDO1*, *KMO*, *kynureninase* (*KYNU*) and *QPRT* as compared the results obtained with plaque homogenates (Fig 1D). Therefore, oxysterols may be the components of plaque microenvironment that contribute to the upregulation of the upstream genes of the KP. Since, oxysterols are established endogenous LXR activators, we exposed primary macrophages to plaque homogenates alone or in combination with the LXR antagonist GSK 2033. *KMO* was the only member of the KP pathway to be clearly downregulated upon LXR blockade (Fig. 1G). The same results were obtained using oxLDLs. Whereas the expression of *IDO-1* in macrophages and dendritic cells is established as well as its key role in these cells^11, 13^, little is known regarding KMO. By re-analyzing the datasets of two independent studies in which single cell RNA seq was performed on mouse atheroma plaque^14, 15^, we observed that the expression levels of *kmo* was very high in macrophages and dendritic cells, similarly to *Ido-1* (Fig. 2A). We confirmed this observation on section of human carotid arteries where we showed that KMO is expressed in plaque macrophages. Interestingly, we noticed that KMO was co-expressed with LXR (Fig. 2B). This corroborates our hypothesis of an LXR-dependent regulation of the expression of *KMO*. To further confirm this assumption, *KMO* expression was assessed in primary macrophages exposed to synthetic LXR agonists (Fig. 2C). Despite a marked increase in *KMO* gene expression upon LXR activation, the screening of human *KMO* gene promoter did not reveal any LXR-responsive element. Therefore, LXR-mediated regulation of *KMO* expression is likely indirect. Consequently, we tested whether the LXR-HIF1α axis could be involved as we showed previously for *IL-1β*, *HK2* or *GLUT1*^3, 4^. Accordingly, we showed that LXR-mediated up-regulation of *KMO* gene expression was reversed in the presence of echinomycin, an HIF-1α antagonist. Using a publically available HIF-1/HIF-2 ChIP-Seq dataset (GSE43109)^16^, we identified a consensus HIF-1-response element (HRE) in human *KMO* gene promoter (Fig. 2E). Then, we performed ATAC-seq experiments on human primary macrophages exposed to atheroma plaque homogenates alone or in combination with the LXR antagonist GSK2033. We were able to show that the exposure to plaque microenvironment affect the binding to the HRE of *KMO* gene promoter in a LXR-dependent manner (Fig. 2F-G).

**Figure 2:**
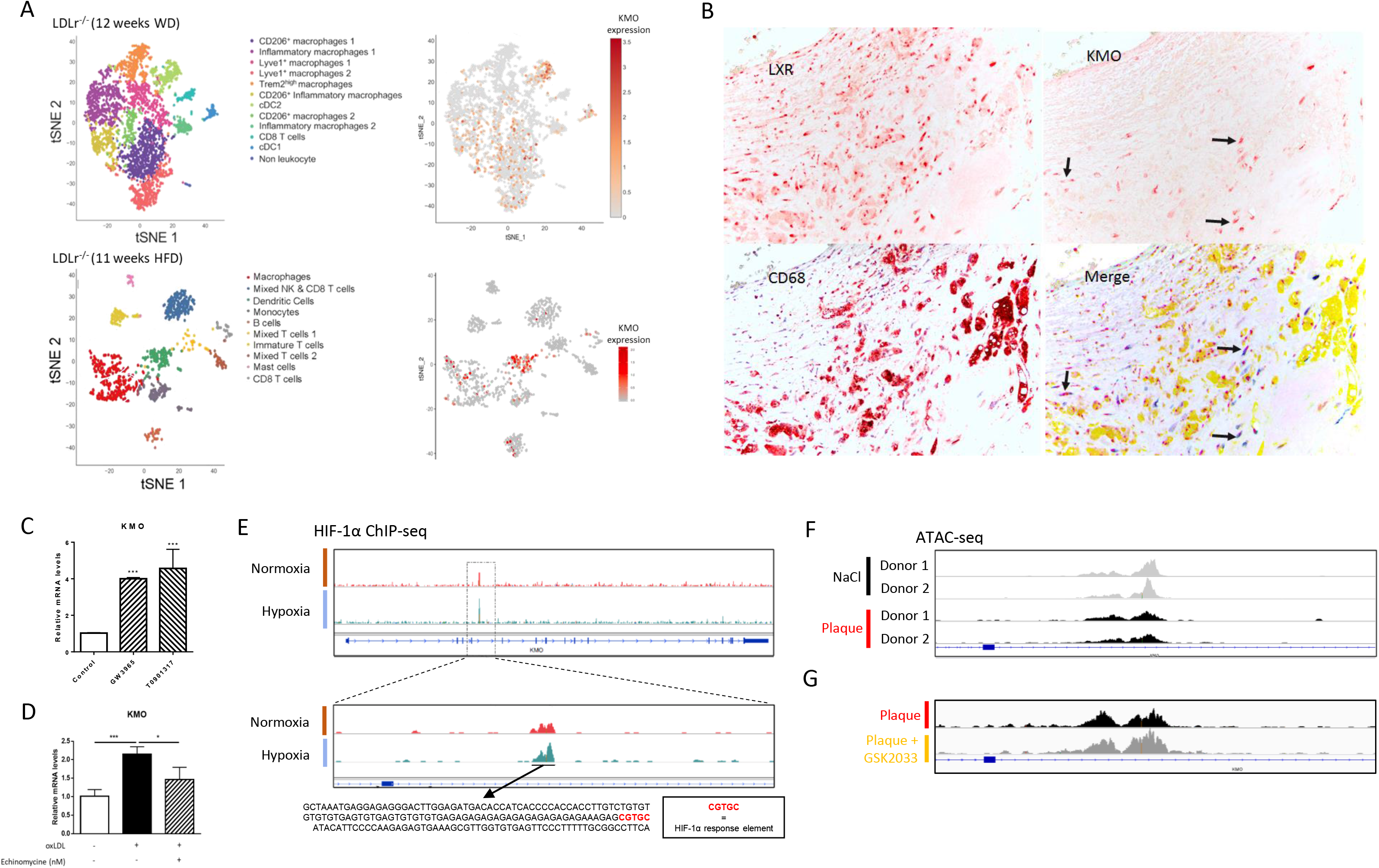
Kynurenine monooxydase expressed in human plaque macrophages is regulated by LXR through an HIF-1α-dependent mechanism. A: KMO expression in mouse atheroma plaque leukocyte subsets. Upper panel, re-analysis of single-cell RNAseq data from from LDLr^-/-^ mice fed for 12 weeks with a western diet (Kim K et al. 2018^14^, GSE116240). CD45^+^ sub-populations are presented on the left and KMO expression in this sub-population of the right part. Lower panel, re-analysis of single-cell RNAseq data from from LDLr^-/-^ mice fed for 11 weeks with a high fat diet (Cochain C et al. 2018^15^, GSE97310). CD45^+^ sub-populations are presented on the left and KMO expression in this sub-population of the right part. B. Representative pictures of sections of human carotid arteries immune-stained for LXR, CD68, KMO. On the merge picture. Yellow color corresponds to CD68 staining, red to LXR staining and blue to KMO staining. Arrows indicate of LXR and KMO colocalization in human plaque macrophages. C. Relative mRNA levels of KMO in primary human macrophages (n=4 healthy donors) treated with the LXR agonists GW3965 (1µM) or T0901317 (100nM) for 24h. Bar graphs represent mean ± s.d.; *** p<0.001 determined by one-way ANOVA corrected for multiple comparison. D. Relative mRNA levels of KMO in primary human macrophages exposed for 24h to oxLDL (50µg/mL) alone or in combination with the HIF-1α inhibitor echinomycine (100nM). Data are representative of 2 independent experiments with different human healthy donors. Bar graphs represent mean ± s.d. * p<0.05; ***: p<0.001 determined by one way ANOVA corrected for multiple comparison. E. Re-analysis of HIF-1α ChIP-seq data from primary human macrophages cultured for 4h in conditions of normoxia or hypoxia (1% O_2_) (GSE43109). Data highlight the presence of HIF-1α response element on KMO gene promoter. F-G. ATAC-seq analysis performed primary human macrophages from healthy donors (F) exposed for 24h to saline or patient atheroma plaque homogenates or (G) to patient atheroma plaque alone or in combination with the LXR antagonist GSK2033. Pictures represent the modulation of HIF-1α binding to KMO gene promoter in the different experimental conditions.

### Plaque microenvironment-induced macrophage kynurenine release induces the expression of adhesion molecules on endothelial cells in AhR-dependent manner

The activation of the KP occurs through the conversion of tryptophan to kynurenine^11^. Thus, we aimed to investigate the influence of the plaque microenvironment on macrophage-mediated kynurenine release. Interestingly, our results revealed a correlation between the extent of kynurenine release and the oxidation degree of LDLs. Specifically, macrophages exposed to highly oxidized LDLs exhibited a higher release of kynurenine. Additionally, we observed that the increased production of kynurenine was associated with a reduction in tryptophan levels in the culture medium of macrophages exposed to oxLDLs (Fig. 3A-B). Exposure to oxLDLs also triggers the release of kynurenic and quinolenic acid but in a less consistent manner compared to what was observed for kynurenine release (Fig. S2). Moreover, through immunostaining experiments using sections of atheroma plaques from human carotid arteries, we successfully detected the presence of kynurenine within the plaque, specifically in the vicinity of macrophages expressing KMO (Fig. 3C). This finding supports the hypothesis of a communication pathway between plaque macrophages and other cellular components of the plaque through the release of kynurenine. Previous studies have described the immunomodulatory properties of KP metabolites, highlighting their role in the interplay between myeloid cells and lymphocytes^14–21^. Building upon this knowledge, we investigated the impact of plaque microenvironment-mediated macrophage kynurenine release on endothelial cells, given their crucial role in the recruitment of blood monocytes to the vicinity of the atheroma plaque.

**Figure 3:**
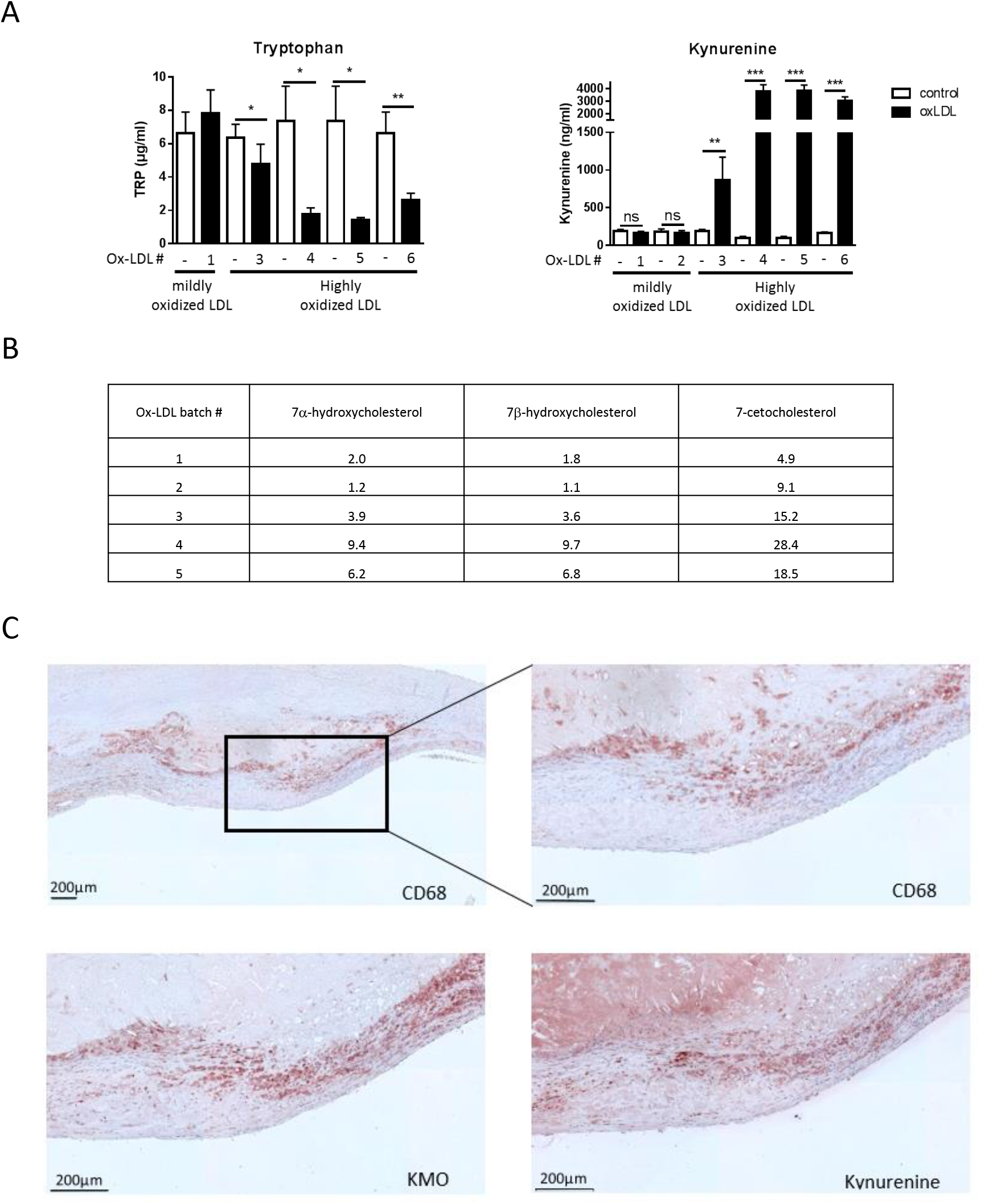
Activation of kynurenine pathway in macrophage by plaque microenvironment leads to kynurenine release. A. Tryptophan (Left) and kynurenine (right) concentrations determined by ELISA in the culture medium of human primary macrophage exposed to different oxLDL batches for 24h (50µg/mL). Data are representative of 5 independent experiments with 5 different healthy donors. Bar graphs represent mean ± s.d. * p<0.05; ** p<0.01; *** p<0.001 determined by Student’s t-test. B: Oxysterol levels in oxLDL batches used in A. Human oxLDL were prepared from LDL isolated from the plasma from 5 healthy donors. The level of oxysterol species is expressed as the ratio of oxysterols to cholesterol in each bacth. Cholesterol 7α-hydroxycholesterol, 7β-hydroxycholesterol and 7-ketocholesterol concentrations were determined by MS/MS and normalized to oxLDL protein content. C. Representative pictures of sections of human carotid arteries stained for CD68, KMO and Kynurenine by immunohistochemistry.

To accomplish this, we exposed human primary endothelial cells (HUVEC) to the culture medium obtained from macrophages exposed to oxLDLs. Subsequently, we confirmed the presence of elevated kynurenine levels in this conditioned medium (Fig. 4A). Intriguingly, the exposure of HUVEC to this kynurenine-enriched conditioned medium was associated with a notable increase in the expression of intercellular adhesion molecule 1 (ICAM1) and vascular cell adhesion molecule 1 (VCAM1) (Fig. 4B). In order to further validate the involvement of kynurenine in this communication between macrophages and endothelial cells, we treated primary human macrophages with two distinct pharmacological inhibitors of KMO. Consequently, a substantial elevation in kynurenine release into the macrophage culture medium was observed, which was subsequently utilized to treat endothelial cells. Importantly, this conditioned medium induced an upregulation in the expression of adhesion molecules within the endothelial cells as well (Fig. 4D). Finally, we exposed HUVEC to escalating concentrations of kynurenine. Notably, kynurenine elicited a dose-dependent upregulation of ICAM-1, both at the protein and mRNA levels, while VCAM-1 expression remained unaffected (Fig. 4E-F). Given that kynurenine has been reported to exhibit the highest AhR agonistic activity among KP metabolites^17^, we investigated whether the induction of endothelial cell adhesion molecules by kynurenine could occur in an AhR-dependent manner. Intriguingly, we observed a dose-dependent reduction in kynurenine-induced upregulation of adhesion molecules in endothelial cells upon treatment with increasing doses of the AhR inhibitor SR-1 (Fig. 4E). Additionally, we evaluated the impact of kynurenine on the activation profile of macrophages exposed to oxLDLs. We observed a significant enhancement of *IL-1β* gene expression, mediated by kynurenine in an AhR-dependent manner, thus indicating its potentiation of oxLDL-induced upregulation (Fig. 4G). Consistent with the findings of Metghalchi *et al.* ^18^, we also observed a reduction in the expression of the anti-inflammatory cytokine *IL-10* in kynurenine-treated macrophages exposed to oxLDL, which was AhR-dependent (Fig. 4G). These results support the notion of a role of kynurenine in promoting an inflammatory profile within the plaque microenvironment. However, further investigations are necessary to fully characterize the macrophage phenotype in this context, as discrepancies in the regulation of other inflammatory cytokines were observed (Fig. 4G).

**Figure 4.**
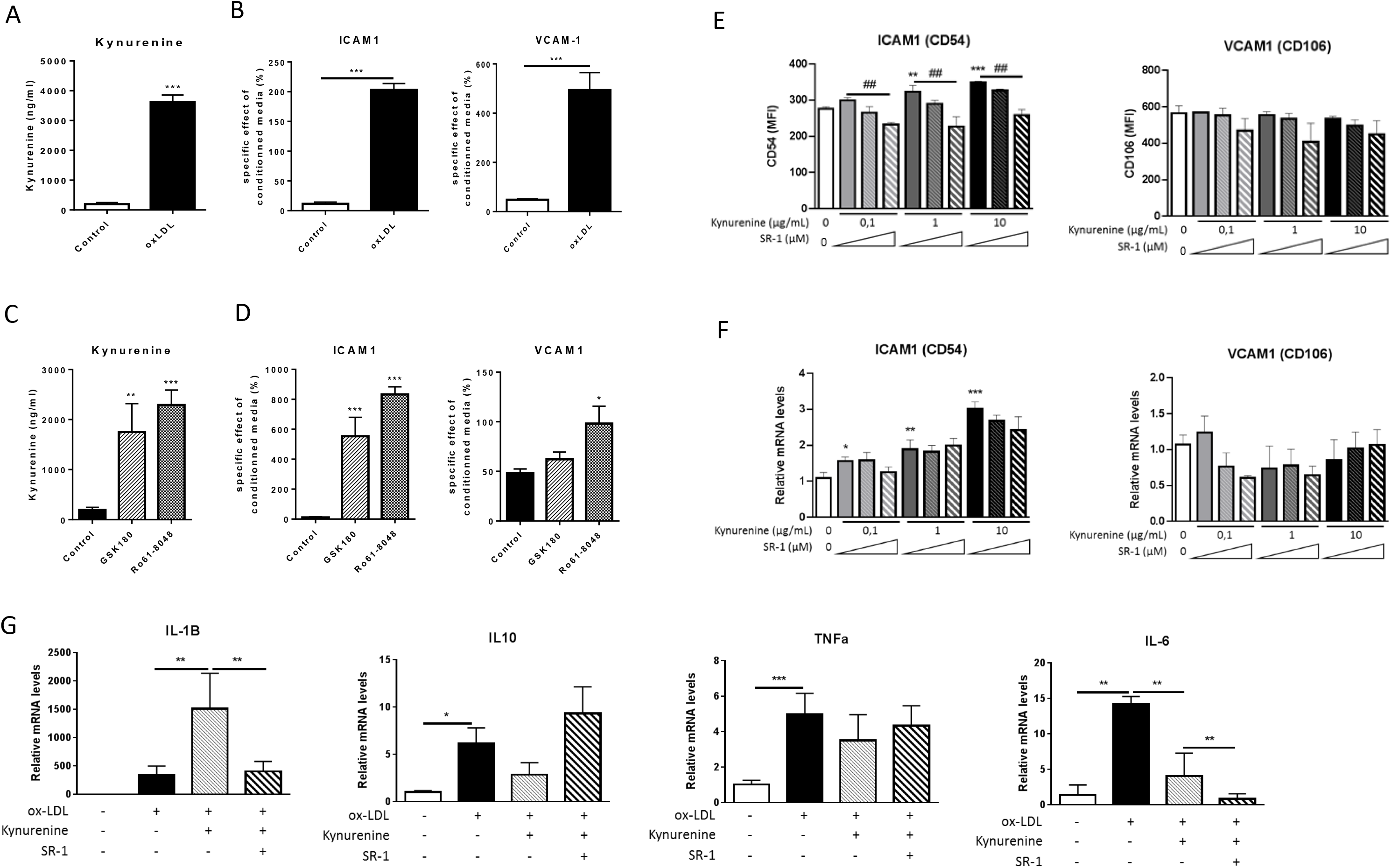
OxLDL-mediated kynurenine release from macrophages induces the expression of endothelial cell adhesion molecules in an AhR-dependent manner. A: Concentrations of kynurenine assessed by ELISA in macrophage-conditioned medium obtained from human primary macrophages treated with oxLDL (50µg/mL) for 24h. B: Expression of adhesion molecules (ICAM1 and VCAM1) measured by flow cytometry in human endothelial cells (HUVEC) exposed to conditioned-media as obtained in A. To neutralize the potential effect of oxLDL, the expression of adhesion molecules in HUVEC cultured with macrophage-conditioned media is expressed in percentage of the expression level of adhesion molecules of HUVEC exposed to the non-conditioned culture medium containing oxLDL. Data are representative of 4 independent experiments. Bar graphs represent mean ± s.d. *** p<0.001 determined by Student’s t-test. Bar graphs represent mean ± s.d. C-D. Experiments performed as in A-B with conditioned-medium obtained from human primary macrophages exposed to the KMO inhibitors GSK180 or Ro61-8048 to induce kynurenine release (C). The normalization was performed as in B but with GSK180 or Ro61-8048 non-conditioned media. Data are representative of 3 independent experiments (C) and 2 independent experiments (D). Bar graphs represent mean ± s.d.; *** p<0.001 determined by one-way ANOVA corrected for multiple comparison. E. Expression of adhesion molecules (ICAM1, VCAM1) measured by flow cytometry on human endothelial cells (HUVEC) after exposure to kynurenine alone or in combination with increasing dose of the AhR inhibitor StemRegenin 1 (SR-1; 0, 0.1, 1µM) for 24h. Controls were treated with vehicle (DMSO). Data are representative of 3 independent experiments with different human donors. Bar graphs represent mean ± s.d. * p<0.05; ** p<0.01; *** p<0.001 (kynurenine versus vehicle), ^#^ p<0.05; ^##^ p<0.01; ^###^ p<0.001 (SR-1 versus kynurenine) comparison between 2 experimental group) determined by one-way ANOVA corrected for multiple comparison. F. Expression of adhesion molecules (ICAM1, VCAM1) measured by qPCR on human endothelial cells (HUVEC) after exposure to kynurenine alone or in combination with increasing dose of the AhR inhibitor StemRegenin 1 (SR-1; 0, 0.1, 1µM) for 24h. Controls were treated with vehicle (DMSO). Data are representative of 3 independent experiments with different human donors. Bar graphs represent mean ± s.d. * p<0.05; ** p<0.01; *** p<0.001 (kynurenine versus vehicle) determined by one-way ANOVA corrected for multiple comparison. G. Relative mRNA levels of IL-1β, IL-10, TNFα and IL-6 in n macrophages exposed to oxLDL (25µg/mL), alone or in combination with kynurenine (1µg/mL) and/or the AhR inhibitor StenRegenin-1 (SR-1; 1µM) for 24h. Controls were exposed to vehicle (DMSO). Data are representative of 2 independent experiments with different human donors. Bar graphs represent mean ± s.d. * p<0.05; ** p<0.01; *** p<0.001 determined by one-way ANOVA corrected for multiple comparison.

### High plasma kynurenine levels are associated with lower extremity arterial disease in atherosclerotic patients

In order to explore the clinical relevance of our in vitro findings and to further understand the role of kynurenine in the onset of atherogenesis, we measured the plasma levels of tryptophan and kynurenine in a cohort 201 atherosclerotic patients enrolled at the university hospital of Dijon (MASCADI cohort: Arachidonic Acid Metabolism in Carotid Stenosis Plaque in Diabetic Patients)^19^. First, on a subset of patients for which sections of carotid arteries with atheroma plaque were available, we checked the level of plaque 27-hydroxycholesterol according to the extent of plaque necrotic core (Fig. S3A). As expected, we observed that a high percentage of necrotic core within atheroma was associated with higher 27-hydroxycholesterol levels in the plaque (Fig. S3A). Interestingly, in accordance with our in vitro experiments with human primary macrophages exposed to atheroma plaque homogenates or oxLDLs, we noticed that patients with the higher levels of plaque 27-hydroxycholesterol presents the highest levels of plasma kynurenine, whereas plasma levels of tryptophan and the kynurenine to tryptophan (kyn/trp) ratio remained unchanged (Fig. 5A). This further support our in vitro data demonstrating that kynurenine production is induced by endogenous LXR ligands in atheroma plaque. In line with these data, we also observed that plasma kynurenine levels were higher in atherosclerotic patients with dyslipidemia (Fig. S3B). More intriguingly, we observed higher plasma levels of kynurenine in patients with lower extremity arterial disease (LEAD) (Fig. 5B). These LEAD patients are at very high risk for the development of carotid atherosclerosis^20^. This explains why we observed higher levels of kynurenine in patients with asymptomatic carotid atherosclerosis (Fig. S3C). Indeed, due to the presence of LEAD (associated with elevated kynureninemia), these patients undergo carotid endarterectomy without the procedure being motivated by myocardial infarction or stroke (referred to as symptomatic carotid atherosclerosis). Noteworthy, the same observation was not made for plasma tryptophan levels and plasma Trp/Kyn ratio (Fig. 5B). Additionally, plasma levels of kynurenine and tryptophan were not associated hyperechogenicity of the atherosclerotic carotid arteries (Fig. 5C). Finally, using multiple regression logistic analysis we observed that in a model including (plasma kynurenine levels, plasma cholesterol levels, smoking, sex, age, BMI, body weight, hypertension and diabetes), plasma kynurenine was independently associated to LEAD (p=0.024). The AUC of the ROC curve obtained with this model was 0.73 which correspond to good predictive value (Fig. 5D). Interestingly, we noticed that taken alone, plasma kynurenine performs better than the classical risk factors sex and smoking status to predict LEAD (Fig. 5E-G). By contrast, the model including plasma kynurenine, sex and smoking status was almost as good the model including all biological variables (Fig. 5H). Finally, this model of multiple logistic regression, we showed that the risk of LEAD was statistically increased in atherosclerotic patients with high plasma levels of kynurenine (Fig. 5I).

**Figure 5.**
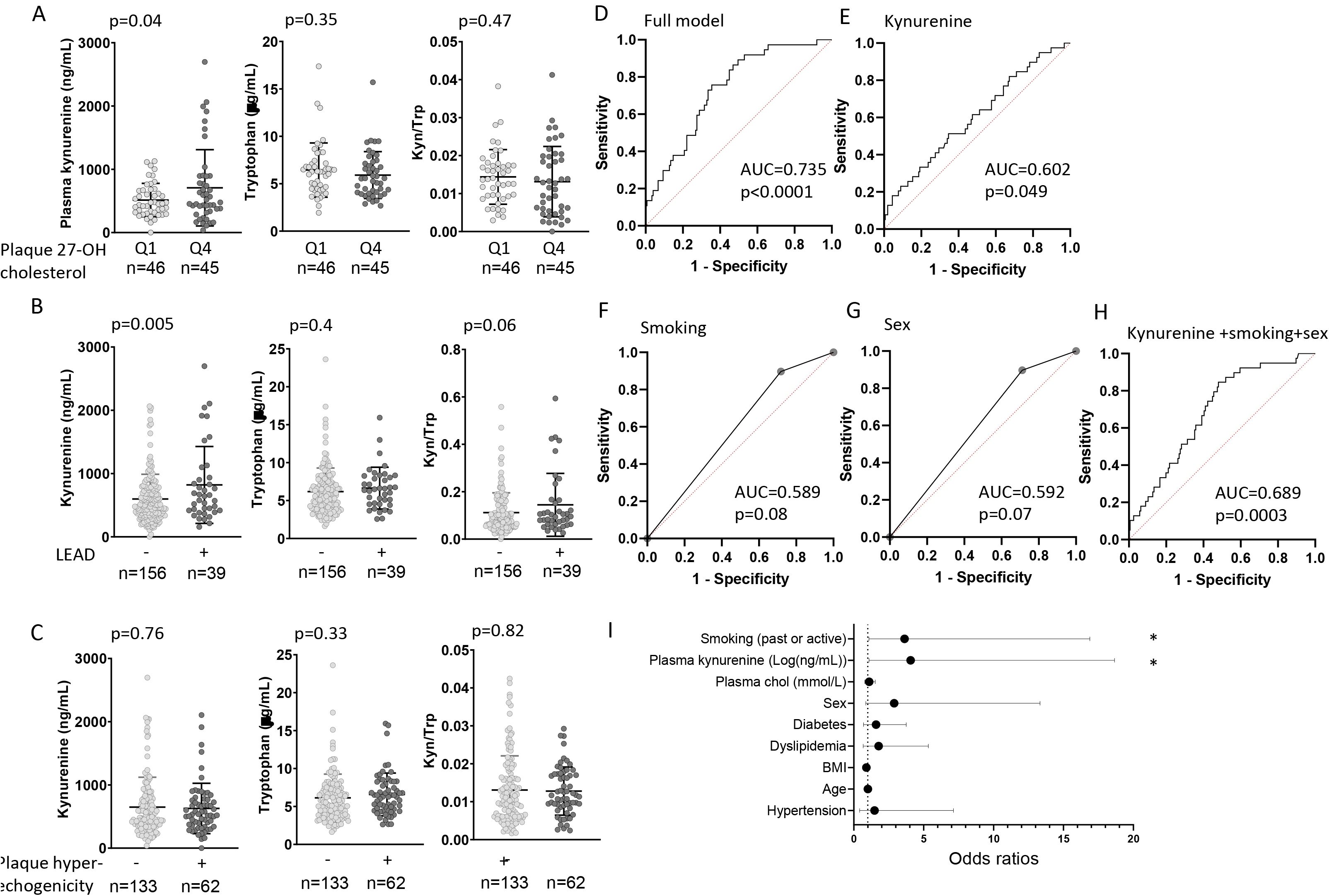
Atherosclerosis is associated with high plasma kynurenine levels in patients. A. Plasma kynurenine levels according to the quartile of 27-hydroxycholesterol levels in plaque carotid arteries from atherosclerotic patients enrolled in MASCADI cohort (n=91). p value determined by Student’s t-test with Welch’s correction. B. Plasma levels of kynurenine, tryptophan and plasma kynurenine to tryptophan (kyn/trp) ratio in atherosclerotic patients from the MASCADI cohort according to lower extremity arterial disease (LEAD) status (n=195). p value was p value determined by Student’s t-test with Welch’s correction. C. Plasma levels of kynurenine, tryptophan and plasma kynurenine to tryptophan ratio in atherosclerotic patients from the MASCADI cohort according to the degree of plaque calcification (n=195). p value determined by Student’s t-test with Welch’s correction. D. Receiver operating characteristic (ROC) curve for the prediction of LEAD in atherosclerotic patients from MASCADI cohort using a model including plasma kynurenine, smoking, age, gender, hypertension, BMI, dyslipidemia, plasma cholesterol, and diabetes. AUC and p value were determined by multiple logistic regression. E-H. ROC, AUC and p value determined as in D with a model including only plasma kynurenine (E), or only smoking status (F), or only sex (G) or the 3 parameters plasma kynurenine, sex and smoking status (H). I. Forest plots showing odds ratio values and 95% confidence intervals for multiple logistic regression analysis of LEAD and targeted biological variables (plasma kynurenine, smoking, age, gender, hypertension, BMI, dyslipidemia, plasma cholesterol, and diabetes). * p<0.05 determined by multiple logistic regression analysis.

## Discussion

Atherosclerosis is a lipid-driven chronic inflammatory disease involving various immune cells that participate in different stages of the disease, including initiation and progression. In the early stages, both innate immune cells (such as dendritic cells, macrophages, and neutrophils) and adaptive immune cells (B and T lymphocytes) contribute to modulating the inflammatory response^21^.

It has now been established that macrophages and dendritic cells undergo significant metabolic reprogramming during their activation. The most well-known change is the shift towards glycolysis, which was initially investigated in vitro through the detection of PAMPs, such as LPS ^22, 23^. There is growing evidence indicating that immune cells also experience metabolic rewiring in the context of atherosclerosis. This phenomenon has been demonstrated in vivo in mice through experimental approaches^14^ and in patients through high-resolution imaging techniques, such as PET scan ^24, 25^. Complementary in vitro investigations, like the plaque homogenate exposure model developed by our laboratory, allow to dissect the underlying molecular mechanisms. Through this approach, we have shown that the LXR-HIFα axis contributes to regulating the inflammatory profile of human macrophages, upregulating IL1β expression, and inducing their metabolic reprogramming towards glycolysis upon exposure to the plaque microenvironment^3, 4^.

Using a similar strategy, we have demonstrated in the present study that the LXR-HIFα axis mediates the effect of the plaque microenvironment on KP up-regulation. We found that exposing macrophages to oxLDL recapitulates this effect. Intriguingly, macrophages release more kynurenine when exposed to oxLDLs that are richer in oxysterols. We made a consistent observation in a cohort of patients with carotid atherosclerosis since plasma kynurenine levels were significantly higher in atherosclerotic patients with the higher oxysterol levels in the plaque. This observation was made with plaque 27-hydroxycholesterol levels but was also true for other oxysterols namely 7α- and 7β-hydroxycholesterol and 7 ketocholesterol (data not shown).

Kynurenine is a product of the KMO enzyme, which plays a pivotal role in the KP as it sits at the crossroads of the QA and KA branches of the pathway. Interestingly, Baumgartner et al., through a transcriptomic approach, demonstrated a deviation of the KP towards the QA branch in the carotid plaques of patients compared to control arteries^26^. Similarly, in our study, we showed a marked upregulation of KMO gene expression. Additionally, we demonstrated that this upregulation of KMO gene expression is induced by plaque microenvironment, especially oxysterols, through LXR-HIFα axis. This observation is significant as it mitigates the protective role of LXR activation in macrophages. Indeed, LXR activation has been linked to enhanced cholesterol efflux^8^. However, unlike in mice, LXR activation in human macrophages does not exhibit a clear anti-inflammatory profile ^4, 5^. This study represents a significant advancement in understanding the connection between LXR signaling and inflammation in human macrophages within the context of atherosclerosis, specifically by examining its impact on the KP.

The KP and its metabolites are closely associated with inflammation. This observation initially stemmed from evidence showing that pro-inflammatory signals, particularly interferons, strongly induce the expression of IDO-1 ^27–30^. This upregulation of IDO-1 is linked to immunomodulation due to Trp depletion or the formation of bioactive metabolites. The shortage of tryptophan associated with IDO-1 induction leads to dysfunction in T lymphocyte differentiation^31, 32^ and an increase in Treg differentiation^33–35^. Moreover, numerous studies, mainly conducted in the context of cancer, have attributed anti-inflammatory properties to several KP metabolites through the activation of AhR signaling. AhR is a ligand-activated transcription factor that exerts potent effects on immune cells and is involved in Treg cell differentiation^36, 37^. Kynurenine has been shown to induce Treg cells through this signaling ^38^. Other KP metabolites, such as xanthurenic acid, kynurinic acid, and cinnabarinic acid, possess immunomodulatory properties and have been demonstrated to activate AhR signaling^20, 39, 40^. In physiopathological settings, kynurenine is likely the most efficient AhR activator ^17^. However, the specific role of kynurenine in atherosclerosis has not been investigated in detail.

In this study, we have demonstrated that plaque microenvironment-mediated macrophage kynurenine release contributes to the increased expression of endothelial adhesion molecules ICAM1 and VCAM1, which play crucial roles in adhesion and recruitment of immune cells to the endothelium, thereby contributing to the progression of atherosclerosis^41^. We have also established that this effect is mediated by AhR signaling, particularly evident for ICAM-1. As ICAM-1 is predominantly expressed on the endothelial surface, it facilitates the transmigration of monocytes into the subendothelial space by interacting with their leukocyte integrin macrophage-1 antigens (Mac-1). This interaction promotes monocyte adhesion, infiltration, and subsequent foam cell formation, ultimately accelerating plaque development^42^.

These observations strongly support the pro-inflammatory role of kynurenine and LXR-mediated activation of the KP. Remarkably, in a cohort of 201 atherosclerotic patients, we found significantly higher plasma kynurenine levels in patients with higher oxysterol content in the plaque. Notably, Metghalchi *et al.* demonstrated that plasma kynurenine levels were substantially reduced in mice grafted with IDO-1^-/-^ bone marrow, suggesting that circulating kynurenine levels likely reflect KP activity in macrophages^18^. In fact, IDO-1 is known to be highly expressed in macrophages^13^. In our study, we further revealed strong KMO expression in macrophages and dendritic cells and highlighted the presence of kynurenine in the vicinity of KMO-expressing macrophages within atheroma plaques from carotid arteries. Although our in vivo data do not provide direct proof, it suggests that kynurenine levels in atherosclerotic patients coincide with KP activation in macrophages triggered by the plaque microenvironment.

In exploring the role of kynurenine in atherosclerotic patients, we observed that plasma kynurenine levels, but not plasma tryptophan levels nor the kyn/trp ratio, were significantly higher in patients with lower extremity arterial disease (LEAD). LEAD results from atherosclerosis in the arteries of the lower limbs and is a risk factor for carotid and coronary atherosclerosis^20^. Intriguingly, by employing a multiple logistic regression model that includes major risk factors of LEAD (smoking, age, gender, hypertension, BMI, dyslipidemia, plasma cholesterol, and diabetes) along with plasma kynurenine levels, we demonstrated that plasma kynurenine is independently associated with LEAD. Furthermore, when using Log-transformed plasma kynurenine levels, we found an odds ratio of 4 for LEAD. Consequently, our data further corroborate previous clinical studies that have emphasized a positive association between IDO activity and the kyn/trp ratio, cardiovascular diseases, and atherosclerosis^43–49^. Although only a few studies have specifically investigated plasma kynurenine levels in atherosclerotic patients, those focused on patients with renal disease revealed positive associations between plasma kynurenine levels, inflammation, and cardiovascular disorders^50^. Interestingly, the same group demonstrated that plasma kynurenine was positively associated with endothelial adhesion molecule markers (soluble ICAM-1 and VCAM-1) and intima-media thickness in atherosclerotic patients^51^. Therefore, our study corroborates these results by showing that intra-plaque kynurenine release may facilitate monocytes recruitment and promote atherosclerosis. Our data are also in line with the study of Metghalchi *et al*. which highlighted that invalidation of IDO-1 prevents atherosclerosis in Ldlr-deficient mice. This was related to a marked decreased Kyn/Trp ratio which led to an increase in IL-10, a major immunoregulatory and atheroprotective cytokine, and to atheroprotection. The authors suggested that the rise in IL-10 was due to the reduction of plasma kynurenine levels, which in turn triggered a decrease in kynurenic acid and abrogated its inhibitory effect on IL-10 production^18^.

Altogether, our data indicate that the activation of KP in macrophages in the context of atheroma plaque is partially mediated by LXR-HIF1α axis and leads to the release of kynurenine. This, in turn, contributes to the exacerbation of both local and peripheral atherosclerosis particularly through the activation of endothelial cells.

### Material and Methods Cell culture

Human peripheral-blood monocytes were obtained from healthy donors with informed consent, provided in accordance with the Declaration of Helsinki by the Etablissement Français du Sang (Besançon, France). Mononuclear cells were isolated by Ficoll gradient centrifugation, and monocyte negative selection was performed via magnetic activated cell sorting using the Monocyte Isolation Kit II (Miltenyi Biotec, Bergish Gladbach, Germany, Cat#130-117-337) according to the manufacturer’s instructions. Monocytes were differentiated into macrophages for 5 days with 100 ng.ml^−1^ of macrophage-colony stimulating factor (CSF1) in RPMI medium supplemented with 10% FBS and maintained in a humidified environment at 37°C with 5% CO2.

Human umbilical cord vein endothelial cells (HUVEC) were prepared from human umbilical cords provided by the maternity unit of Dijon University Hospital. Cells were dissociated from the umbilical cord vein with collagenase A 0.2% (Roche Diagnostics), harvested, and cultured in Endothelial Cell Growth Basal Medium 2 (EBM-2; Lonza) supplemented with 2% FBS, bovine brain extract, ascorbic acid, human epidermal growth factor, gentamicin sulfate and hydrocortisone (EBM SingleQuots kit; Lonza). HUVEC were cryopreserved at passage 1 and used for experiments between passage 3 and 5.

### Cell treatments

#### Preparation of atheroma plaque homogenates

Atheroma plaque samples were obtained from patients undergoing carotid endarterectomy at the Department of Cardiovascular Surgery at the University Hospital of Dijon (Burgundy, France) (MASCADI, NCT03202823)^19^. All patients provided an informed consent and the study was approved by the local ethic committee (CPP 17.02.05). Lipid rich cores of plaque samples were carefully dissected under aseptic conditions and were homogenized with three volumes of cold NaCl 150 mmol.L^−1^ and briefly sonicated. Macrophages were treated with whole homogenate for 24 or 48 h, as indicated in the figure legends, alone or in combination with LXR antagonist, KMO inhibitors or HIF-1α inhibitor (see below). Experiments were performed with plaque homogenates from different donors in order to take inter-individual variability into account.

#### Preparation of oxidized LDLs (oxLDLs)

Low-density lipoproteins (LDLs) were isolated by sequential ultracentrifugation from human plasma, obtained from healthy donors with informed consent, provided in accordance with the Declaration of Helsinki by the Etablissement Français du Sang (Besançon, France). First, plasma was centrifuged at 190000g for 20h, after density adjustment to 1.019, to separate very low-density lipoproteins and chylomicron from LDL and high-density lipoproteins. LDL were then separated from HDL with a second ultracentrifugation at 220000g for 22h, by adjusting fraction density to 1.063. All ultracentrigufation steps were performed with a 70.Ti rotor in an Optima XPN-90 ultracentrifuge (Beckman, Palo Alto, CA). Density adjustment was achieved by adding potassium bromide to the different fractions. After isolation, LDLs were dialyzed agains 0.9% NaCl and oxidized with 5µM of copper sulfate, overnight at 37°C. EDTA to a final concentration of 200µM was added to the reaction mix in order to stop the oxidation reaction. Residual copper sulfate and potassium bromide were removed from the oxLDLs solution by extensive dialysis. LDL oxidation was confirmed by agarose electrophoresis with Soudan black staining to assess the shift in electrostatic charge (Biotec Fischer, Reiskirchen, Germany).

#### Primary macrophages

After differentiation, macrophages were treated with oxLDL at indicated concentrations (25 or 50µg/mL) or atheroma plaque homogenates (1%v/v) alone or in combination with LXR antagonist GSK2033 (1 to 5µM, Sigma-Aldrich); or the KMO inhibitors GSK180 (10 to 100nM, Sigma-Aldrich) or Ro61-8048 (1 to 10µM) (Sigma-Aldrich); or the HIF-1α inhibitor echinomycin (5µM, Sigma-Aldrich).

#### Human umbilical cord vein endothelial cells (HUVEC)

For experiments with contioned-media from treated primary macrophages, after treatment, cell culture supernatants from macrophages were harvested. The supernatant was centrifuged at 300g for 8 min to pellet cell debris. HUVEC were cultured then for an additional 24h period with complete EBM-2 supplemented with macrophage conditioned-medium (v/v) alone or in combination with kynurenine (0.1ng/mL to 10µg/mL, Sigma-Aldrich) and/or the AhR inhibitor StemRegenin 1 (1 to 10µM, Biotechne).

### Gene expression analysis

RNA isolation and qPCR Total RNA was extracted using TriZol Reagent (Invitrogen) or RNeasy Mini Kit (Qiagen, Hilden, Germany, Cat#74106) according to the manufacturer’s instructions; 100 ng of total RNA was reverse transcribed using M-MLV Reverse Transcriptase (Cat#28025013), random primers (Cat#48190011), and RNaseOUT inhibitor (Cat#10777019) (Invitrogen, Carlsbad, California, USA). cDNA obtained were quantified by real time PCR using PowerUp™ SYBR™ Green Master Mix (Applied Biosystems) and using a StepOne Plus Real-Time PCR System (Applied Biosystems, California, USA). ΔΔCt method was used to determine the relative mRNA levels and Ct were normalized using 36B4 mRNA or 18s rRNA levels. Values are set at 1 in the control group. Primer sequences are available on request.

### RNA sequencing

Macrophages from three different donors, exposed to distinct plaque homogenates alone or in combination with GSK2033, were used for RNA sequencing. Total RNA were prepared by using Qiagen RNA Easy Kit (Qiagen, Hilden, Germany, Cat#74106). mRNA purification from total RNA and library preparation were performed with the NEBNext Ultra RNA Library Preparation Kit with poly-A selection (New England Biolabs, Ipswich, Massachusetts, USA, Cat#E7530S) according to the manufacturer’s protocols. Libraries were sequenced with 2×150bp paired-end reads on an Illumina HiSeq. Sequence reads were trimmed to remove possible adapter sequences and nucleotides with poor quality using Trimmomatic v.0.36. The trimmed reads were mapped to the *Homo sapiens* GRCh38 reference genome using the STAR aligner v.2.5.2b. Unique gene hit counts were calculated by using featureCounts from the Subread package v.1.5.2. Only unique reads that fell within exon regions were counted. After extraction, the gene hit counts table was used for downstream differential expression analysis. Using DESeq2, a comparison of gene expression between the groups of samples was performed. The data discussed in this publication have been deposited in NCBI’s Gene Expression Omnibus (GSE125126).

### Flow cytometry

HUVEC were dissociated from culture plates with Trypsine-EDTA (0,05%; Thermofisher). Cells were stained for 30 min with fixable viability staining 510 (BD Biosciences), followed by a staining with specific anti-human CD54-PE (BD Biosciences, Cat#555511) and anti-human CD106-APC (Biolegend, Cat#B316538) antibodies. Flow cytometry analysis were performed with a BD LSR II flow cytometer (BD Biosciences). Flow cytometry data were analyzed with FlowJo v10 software (BD Biosciences).

### Immunohistological analysis of human carotid plaques

The plaque samples were fixed in neutral buffered formalin, dehydrated in graded alcohols, cleared in xylene, and embedded in paraffin. Slices of 5 μm thick were deposited on slide to achieve a haematoxylin–eosin staining and immunohistochemical analysis. Automated immunohistochemistry (Leica BondMax) was performed using standard procedures on serial sections of plaques.

The following antibodies were used: mouse monoclonal anti-LXRα/NR1H3 antibody (OTI1A5) (1/200, Novus Biologicals, Cat# NBP2-46220), mouse monoclonal anti-CD68 antibody (1/200, Invitrogen, Cat#14-0688-82), KMO (1/250; Sigma Cat#HPA031115), kynurenine (1/25; ImmuSmol, Cat#14IS003), kynurenic acid (1/50; ImmuSmol, Cat#IS010) and quinolinic acid (1/50; ImmuSmol, Cat#1IS002). Revelation was made using ImmPress anti-Mouse kit (Vector Labs, Cat#MP2400) for anti-CD68, anti-kynurenine, anti-kynurenic acid and anti-quinolinic acid antibodies and ImmPress anti-Rabbit kit (Vector Labs, Cat#MP7401) for the anti-KMO antibody. Chromogenic revelation was performed using AEC (Vector Labs - SK-4200).

For multiple labelling experiments, tissue sections were incubated in 3% hydrogen peroxide for 15 min and in 3% BSA for 20 min before adding the primary antibody, followed by Polymer-HRP detection system (Leica, Novolink Polymer Detection Systems), and chromogenic revelation was performed using AEC (Vector, SK-4200). Tissue sections were then mounted with aqueous mounting medium (Aquatex®, Merck #108562) and scanned for digital imaging (Zeiss Axioscope A1 microscope equipped with a Jenoptic Gryphax camera). After scanning, slide coverslips were directly removed in water, and tissue sections were destained in ethanol. Then, the slides were subjected to the next round of staining with the next primary antibody as described (Remark et al., 2016). Individual images were converted to 8-bit grayscale files with ImageJ and merged to RGB files. Tissue sections were scanned for digital imaging with a Zeiss Axioscope A1 microscope equipped with a Jenoptic Gryphax camera.

#### Tryptophan, kynurenine, kynurenic acid and quinolenic acid quantification

Tryptophan, kynurenine, kynurenic acid and quinolenic acid concentration were determined in plasma (tryptophan, kynurenine) and cell culture supernatants (tryptophan, kynurenine, kynurenic acid and quinolenic acid) by enzyme linked immuno-sorbent assay (ELISA). L-Kynurenine, L-Tryptophan, kynurenic acid and quinolenic acid ELISA kit were all from Immusmol and were used following manufacturer’s instructions.

### Bioinformatics analysis & ATAC seq

#### Single cell RNA sequencing data analysis

Single cell RNA sequencing data were obtained from the Gene Expression Omnibus database. 2 different datasets were analyzed. GSE116240 represents data obtained in LDLr^-/-^ mice placed on Western diet for 12 weeks. CD45^+^ cells isolated from mice aorta were analyzed. GSE97310 represents data obtained in LDLr-/-mice placed with high fat diet for 11 weeks. CD45+ cells isolated from mice aorta were analyzed. Cells were represented with t-SNE plots. KMO expression in different cells was obtained with Deseq2 R/Bioconductor package. **ChIP-seq data analysis**. ChIP-seq data were obtained from the Gene Expression Omnibus database (GSE43109). Data were analyzed using Integrative Genomics Viewer (Robinson JT et al. 2011). Hg38 reference genome was used to identify genes and regions of interest.

#### ATAC-seq

ATAC-seq analysis were performed by Active Motif company (Waterloo, Belgium) on 10^5^ cells per sample. After 48h of treatment with atheroma plaque homogenates alone or in combination with the LXR antagonist GSK2033, cells were harvested, stored at −80°C and transfer to Active Motif for tagmentation, library preparation and sequencing. Filtered reads were aligned to the Hg38 reference genome. Open chromatin regions were identified with HOMER. Peaks were analyzed and visualized using Integrative Genomics Viewer (IGV) ^52^.

#### Promoter analysis

Identification of predicted DR4-type response element within LXR peak regions or hypoxia-response element within HIF1a peak regions were performed using the nuclear hormone receptor (NHR)-scan web tool available at http://www.cisreg.ca/cgi-bin/NHR-scan/nhr_scan.cgi. Motifs identified were compared against the database HOMOCO (v11 full) of known motifs using TOMTOM Motif Comparison Tool available at http://meme-suite.org/tools/tomtom

#### Oxysterol quantitation by gas chromatography-mass spectrometry (GC–MS)

oxLDL sample were mixed with NaCl 0.9% and saponified for 45 min at 56°C with 60 μl of potassium hydroxide 10 mol.L^−1^ and 1.2 ml of ethanol-BHT (50 mg.L−1) containing 20 μl of standard-intern mix (μg per sample: 0.2 (25-OH-cholesterol-d6); 0.4 (27-OH-cholesterol-d6); 0.5 (7α-OH-cholesterol-d7); 0.5 (7β-OH-cholesterol-d7); 1 (7-keto-cholesterol-d7)). Sterols were extracted with 5 ml of hexane and 1 ml of water. After evaporation of the organic phase, sterols were derivatized with 100 μl of a mixture of bis (trimethylsilyl)trifluoroacetamide/trimethylchlorosilane 4/1 v/v for 1 h at 80°C. After evaporation of the silylating reagent, 100 μl of hexane were added. Trimethylsilylethers of sterols analysis was performed by GC–MS in a 7890A gas chromatograph coupled with a 5975C Mass Detector (Agilent Technologies). Separation was achieved on a HP-5MS 30 m × 250 μm column (Agilent Technologies, Santa-Clara, USA) using helium as carrier gas and the following GC conditions: injection at 250°C using the pulsed split, oven temperature program: initial temperature 150°C up to 280°C at a rate of 15°C.min^−1^, up to 290°C at a rate of 1°C.min−1 for 2 min. The MSD was set up as follow: EI at 70 eV mode, source temperature at 230°C. Data were acquired in SIM mode using following quantitation ions (m/z): 131.1 for 25-OH-cholesterol; 137.1 for 25-OH-cholesterol d6; 145.1 for 24(S)-OH-cholesterol; 151.1 for 24(S)-OH-cholesterol-d6; 368.3 for cholesterol; 375.3 for cholesterol-d7; 417.4 for 27-OH-cholesterol; 423.4 for 27-OH cholestrol-d6; 456.4 for 7α–7β cholesterol; 463.5 for 7α–7β cholesterol-d7; 472.4 for 7-keto-cholesterol; 479.4 for 7-keto-cholesterol d7. A calibration curve was performed for cholesterol with standards.

### Human studies

The MASCADI^19^ protocol was reviewed and approved by the regional Ethics Committee (Comité de Protection des Personnes Est, Dijon, France CPP Est III, CHRU Nancy, N° 2017-A00022-51) and recorded on ClinicalTrials.gov (clinical registration number: NCT03202823). Two-hundred-and-one patients were enrolled. Patients enrolled in the present study were admitted at the Department of Cardiovascular Surgery at the University Hospital of Dijon (Burgundy, France) for the surgical treatment of an atheromatous internal carotid artery (ICA) stenosis whether symptomatic or not. A stenosis was considered symptomatic if stroke or transient ischemic attack (TIA) attributed to the ICA lesion occurred within the previous 6 months before intervention. According to the trial category (Interventional research that does not involve drugs or non-CE-marked medical devices and that involves only minimal risks and constraints for the patient) and the ethics committee, all patients received written information note on the study. Oral consent was obtained from the patient and an attestation of the patient’s oral consent was signed by the investigator physician and countersigned by the patient, according to the French law in this type of trial. Blood serum samples and EDTA total blood samples were collected in parallel on the day of surgery. For all patients, carotid endarterectomy (CEA) was performed within the carotid bulb, with en-bloc removal of the entire plaque, and resulting samples were frozen at −80 °C prior to lipidomic analysis.

Clinical data, imaging plaque characteristics, and biological analysis Demographics, vascular risk factors (hypertension, active smoking, type 2 diabetes, dyslipidemia with detailed treatment, chronic renal insufficiency) and symptomatic stenosis were collected. Preoperative plaque imaging assessment included for all patients doppler ultrasound evaluating plaque echolucency and degree of stenosis according to NASCET measures, and CT angiographies (CTA) describing the presence of plaque ulceration and calcification. Glycated hemoglobin A1c (HbA1c) was measured by high pressure liquid chromatography on a Tosoh G8 analyzer (Tosoh Bioscience, Tokyo, Japan), C reactive protein (CRP), troponin I, creatinine, total cholesterol, HDL cholesterol and triglycerides were determined on a Dimension Vista analyzer using dedicated reagents (Siemens). Serum IL-1β levels were determined using a high-sensitivity enzyme-linked immunosorbent assay (ThermoFisher Scientific) with a limit of quantification (LOQ) of 0.05 pg/mL^19^. Quantification of plaque sterols and oxysterols was performed by gas chromatography mass spectrometry (GC–MS).

### Statistical analysis

The number of experiments for each panel is n = 3 to 5 independent experiments with individual donors or individual plaques samples. RNA seq analysis was performed with three independent donors. These transcriptomic data were replicated by qPCR in n = 5 independent donors. The average of the technical replicates was used when combining data from independent donors. No outliers were excluded. In QCPR experiments, the mean of ΔC(t) values from the control group (untreated samples) was used as a reference in the ΔΔC(t) calculation. Data are expressed as ‘Relative mRNA levels (fold mean of the control)’. Statistical differences were analyzed with GraphPad Prism 8. Comparisons of two groups were calculated with unpaired Student’s *t*-test with Welch’s correction. For comparisons with more than two groups, Brown-Forsythe and Welch ANOVA tests method were used. Correction for multiple comparisons was performed with the two-stage set-up method of Benjamini, Krieger and Yekutieli. Correlations were analyzed with Spearman’s rank correlation. A *P* value of less than 0.05 was considered statistically significant. Results are presented as mean ± sd.

In human studies, association between biological parameters and disorders was assessed by multiple regression analysis. Receiver Operating Characteristic (ROC) analysis was utilized to evaluate the predictive accuracy of the identified parameters. Odds ratios were computed to quantify the magnitude of the effect of each parameter. All experimental values were obtained from the measurement of distinct samples and non-repeated measures of the same sample. The number of replicates is indicated in the Figure legends. In vitro experiments were repeated at least two times.

### General methods statements

Operators were not blinded to the experimental treatments; however, data analysis was performed semi-blinded by an independent analyst.

## Acknowledgments

For their excellent technical assistance, the authors thank Hélène Choubley, Victoria Bergas and Jean-Paul Pais de Barros, from the lipidomic facility of the university of burgundy (LAP); Nicolas Pernet, Serge Monier, Amandine Bataille and Anabelle Legrand from the Imaging and flow cytometry facility (Imaflow) of the University of Burgundy.

